# Biomechanical study of cellulose scaffolds for bone tissue engineering *in vivo* and *in vitro*

**DOI:** 10.1101/2021.07.07.451476

**Authors:** Maxime Leblanc Latour, Maryam Tarar, Ryan J. Hickey, Charles M. Cuerrier, Isabelle Catelas, Andrew E. Pelling

## Abstract

Plant-derived cellulose biomaterials have recently been utilized in several tissue engineering applications. These naturally-derived cellulose scaffolds have been shown to be highly biocompatible *in vivo*, possess structural features of relevance to several tissues, and support mammalian cell invasion and proliferation. Recent work utilizing decellularized apple hypanthium tissue has shown that it possesses a pore size similar to trabecular bone and can successfully host osteogenic differentiation. In the present study, we further examined the potential of apple-derived cellulose scaffolds for bone tissue engineering (BTE) and analyzed their mechanical properties *in vitro* and *in vivo*. MC3T3-E1 pre-osteoblasts were seeded in cellulose scaffolds. Following chemically-induced osteogenic differentiation, scaffolds were evaluated for mineralization and for their mechanical properties. Alkaline phosphatase and Alizarin Red staining confirmed the osteogenic potential of the scaffolds. Histological analysis of the constructs revealed cell invasion and mineralization throughout the constructs. Furthermore, scanning electron microscopy demonstrated the presence of mineral aggregates on the scaffolds after culture in differentiation medium, and energy-dispersive spectroscopy confirmed the presence of phosphate and calcium. However, although the Young’s modulus significantly increased after cell differentiation, it remained lower than that of healthy bone tissue. Interestingly, mechanical assessment of acellular scaffolds implanted in rat calvaria defects for 8 weeks revealed that the force required to push out the scaffolds from the surrounding bone was similar to that of native calvarial bone. In addition, cell infiltration and extracellular matrix deposition were visible within the implanted scaffolds. Overall, our results confirm that plant-derived cellulose is a promising candidate for BTE applications. However, the discrepancy in mechanical properties between the mineralized scaffolds and healthy bone tissue may limit their use to low load-bearing applications. Further structural re-engineering and optimization to improve the mechanical properties may be required for load-bearing applications.

## 1. INTRODUCTION

Large bone defects caused by injury or disease often require biomaterial grafts to completely regenerate (1). Current techniques designed to enhance bone tissue regeneration commonly employ autologous, allogeneic, xenogeneic, or synthetic grafts (2). Autologous bone grafting, for which the material is derived from the patient, is considered the “gold standard” grafting practice in large bone defect repairs, but there are several drawbacks including size and shape limitations, tissue availability, and sampling site morbidity (3). In addition, autologous grafting procedures are prone to infections, subsequent fractures, hematoma formation at the sampling or repaired site, and post-operative pain (4). Bone tissue engineering (BTE) provides a potential alternative to traditional bone grafting methods (5). It combines the use of structural biomaterials and cells to create new functional bone tissue. The biomaterials used for BTE should be designed with a macroporous architecture, surface chemistry for cell attachment, and mechanical properties similar to those of the native bone (6). Previous studies have shown that the optimal pore size for biomaterials used for BTE is approximately 100-200 μm (7), and the optimal elastic modulus is 0.1 to 20 GPa depending on the grafting site (8). Moreover, the porosity and pore interconnectivity are two important factors that affect cell migration, nutrient diffusion, and angiogenesis (8).

BTE has shown promising results with a diverse set of biomaterials developed as alternatives to bone grafts. These biomaterials include osteoinductive materials, hybrid materials, and advanced hydrogels (8). Osteoinductive materials induce the formation of *de novo* bone structure. Hybrid materials are made of synthetic and/or natural polymers (8). Advanced hydrogels mimic the extracellular matrix (ECM) and deliver the required bioactive agents to promote bone tissue integration (8). Hydroxyapatite is a traditional material choice for BTE due to its biocompatibility and composition (9). Another type of biomaterial for BTE is bioactive glass, which stimulates specific cell responses to activate genes necessary for osteogenesis (10,11). Biodegradable polymers such as poly(glycolic acid) and poly(lactic acid) are also widely used for BTE (12). Finally, natural (or naturally-derived) polymers such as chitosan, chitin, and bacterial cellulose have also shown promising results for BTE (13). Although these polymers, either natural or synthetic, show some potential for BTE, extensive protocols are usually required to obtain a functional biomaterial with a desired macrostructure. Conversely, native macroscopic cellulose structures can be derived from various plants. In a previous study, our group demonstrated that following a simple surfactant treatment, cellulose-based scaffolds derived from plants can be used as a material for various tissue reconstruction, taking advantage of the native structure of the plant (14). Furthermore, these biomaterials can be used for *in vitro* mammalian cell culture (14), are biocompatible, and can become spontaneously vascularized subcutaneously (14–16). Our group and others have shown that these biomaterials can be sourced from specific plants according to the intended application (14–18). For instance, the vascular structure from plant stems and leaves displays a similar structure to the one found in animal tissues (18). Plant-derived cellulose scaffolds can also easily be carved into specific shapes and treated to alter their surface biochemistry (16). In a recent study, we included a salt buffer in the decellularization process, which resulted in an increase in cell attachment, both *in vitro* and *in vivo* (16). In the same study, we showed that plant-derived cellulose can be used in composite biomaterials by casting hydrogels onto the scaffold surface. In recent studies, functionalization of plant-derived scaffolds has proven to improve their efficiency (17). For example, Fontana et al. (2017) showed that RGD-coated decellularized stems support the adhesion of human dermal fibroblasts as opposed to non-coated stems. Furthermore, the authors also showed that decellularized plant stems can be artificially mineralized in modified simulated body fluid. More recently, Lee et al. (2019) employed plant scaffolds to generate bone-like tissues *in vitro*. After assessing a variety of plants, the authors found that apple-derived scaffolds were the most suitable for the culture and differentiation of human induced pluripotent stem cells (hIPSCs). They also suggested that structural and mechanical properties of the apple-derived scaffolds play a key role. Apple-derived scaffolds were the first plant-derived scaffold employed in tissue engineering applications and have been demonstrated to have a very similar architecture to bone, especially with regards to their interconnected pores ranging from 100 to 200 μm in diameter (14,19).

In the present study, we further examined the potential of apple-derived cellulose scaffolds for BTE and analyzed their mechanical properties *in vitro* and *in vivo*. While the potential use of apple-derived scaffolds for BTE applications has been studied (19), the mechanical properties of the scaffolds have not been investigated systematically. Results of the present study show that scaffolds seeded with MC3T3-E1 and cultured in differentiation medium have a Young’s modulus of 192.0 ± 16.6 kPa, which is significantly higher than those of cell-seeded scaffolds cultured in non-differentiation medium (24.1 ± 8.8 kPa) and acellular scaffolds (31.6 ± 4.8 kPa). Nevertheless, the Young’s modulus of healthy bone tissue is typically in the range of 0.1 to 2 GPa for trabecular bone and between 15 and 20 GPa for cortical bone (8). Moreover, after implantation for 8 weeks in a rodent calvarial defect model, the cell-seeded scaffolds were integrated into the surrounding bone, requiring a force of 113.6±18.2 N to push out the scaffolds from the surrounding bone, similar to values reported in previous studies of cortical bone displacement (20). While these results are very promising, especially for non load-bearing applications, apple-derived cellulose scaffolds still lack the appropriate mechanical properties to match the surrounding bone tissue at an implant site. Further development is therefore required for these scaffolds to reach their full potential.

## 2. MATERIALS AND METHODS

### 2.1. Scaffold preparation

Samples were prepared following established methods (16). Briefly, McIntosh apples (Canada Fancy) were cut in 8 mm-thick slices with a mandolin slider. The hypanthium tissue of the apple slices was cut into squares of 5 mm by 5 mm. The square tissues were decellularized in 0.1% sodium dodecyl sulfate (SDS, Fisher Scientific, Fair Lawn, NJ) for two days. Decellularized samples were then washed in deionized water, followed by an overnight incubation at room temperature in 100 mM CaCl_2_ to remove the remaining surfactant. The samples were subsequently sterilized with 70% ethanol for 30 min, washed with deionized water, and placed in a 24-well culture plate prior to cell seeding.

### 2.2. Cell culture and scaffold seeding

MC3T3-E1 Subclone 4 cells (ATCC® CRL-2593™, Manassas, VA) were maintained at 37°C in a humidified atmosphere of 95% air and 5% CO_2_. The cells were cultured in culture medium made of Minimum Essential Medium (α-MEM; ThermoFisher, Waltham, MA) supplemented with 10% fetal bovine serum (FBS; Hyclone Laboratories Inc., Logan, UT) and 1% penicillin/streptomycin (Hyclone Laboratories Inc.), before being trypsinized once they reached 80% confluency. They were then resuspended in culture medium.

A 40 μL aliquot of cell suspension containing 10^6^ cells was pipetted on the scaffolds. The cells were left to adhere for 1h in cell culture conditions (i.e., at 37°C in a humidified atmosphere of 95% air and 5% CO_2_). Subsequently, 2 mL of culture medium were added to each culture well. Culture medium was changed every 2-3 days, for 14 days. After this incubation, differentiation of MC3T3-E1 cells was induced by adding 50 μg/mL ascorbic acid and 4 mM sodium phosphate to the culture medium (differentiation medium). Differentiation medium was changed every 3-4 days, for 4 weeks. Scaffolds in non-differentiation culture medium (without the supplements to induce differentiation) were incubated for the same duration, with the same medium change schedule, and served as a negative control. All subsequent analyses were conducted at the end of the 4-week incubation. Finally, the cell-seeded scaffolds were imaged after the 4-week incubation using a 12-megapixel digital camera and compared to scaffolds with no seeded cells.

### 2.3. Pore size measurements and cell distribution analysis using confocal laser scanning microscopy

To measure the scaffold pore size, decellularized apple scaffolds (prior to MC3T3-E1 cell seeding) (n=3) were thoroughly washed with phosphate buffered saline (PBS; ThermoFisher) and incubated in 1 mL of 10% (v/v) Calcofluor White solution (Sigma-Aldrich, St. Louis, MO) for 25 min in the dark and at room temperature for staining. Subsequently, the scaffolds were washed with PBS and imaged with a high-speed resonant confocal laser scanning microscope (Nikon Ti-E A1-R; Nikon, Mississauga, ON). ImageJ software was used to process and analyze the confocal images. Briefly, maximum projections in the Z axis were created and the Find Edges function was used to highlight the edge of the pores. A total of 54 pores were analyzed (6 pores in 3 randomly selected areas per scaffold, with 3 scaffolds). Pores were manually traced using the freehand selection tool in ImageJ. The selections were fit as an ellipse to output the major axis length.

To analyze MC3T3-E1 cell distribution, cell-seeded scaffolds cultured in non-differentiation or differentiation medium (n=3 for each condition) were washed with PBS (without Ca^2+^ and Mg^2+^) and fixed with 4% paraformaldehyde for 10 min. They were then washed with deionized water prior to permeabilizing the cells with a Triton-X 100 solution (ThermoFisher) for 5 min, and washed again with PBS. Staining of the scaffolds was performed as previously described (14–16). Briefly, the scaffolds were incubated in 1% periodic acid (Sigma-Aldrich) for 40 min. After rinsing with deionized water, they were incubated for 2h in the dark and at room temperature in 100 mM sodium metabisulphite (Sigma-Aldrich) and 0.15 M hydrochloric acid (ThermoFisher) supplemented with 100 μg/mL propidium iodide (Invitrogen, Carlsbad, CA). Finally, they were washed with PBS, stained with 5 mg/mL DAPI (ThermoFisher) for 10 min in the dark, washed again, and stored in PBS prior to imaging. The surfaces of the cell-seeded scaffolds were imaged with the high-speed resonant confocal laser scanning microscope (Nikon Ti-E A1-R). ImageJ software was used to process the confocal images and create a maximum projection in the Z axis for image analysis.

### 2.4. Alkaline phosphatase and calcium deposition

Before staining with either 5-bromo-4-chloro-3′-indolyphosphate and nitro-blue tetrazolium (BCIP/NBT, Sigma-Aldrich) for alkaline phosphatase (ALP) activity or Alizarin Red S (ARS, Sigma-Aldrich) for calcium deposition, scaffolds were washed three times with PBS (without Ca^2+^ and Mg^2+^) (Hyclone Laboratories Inc.) and fixed with 10% neutral buffered formalin for 30 min.

BCIP/NBT staining solution was prepared by dissolving one BCIP/NBT tablet (Sigma-Aldrich, Cat. No. B5655) in 10 mL of deionized water. After fixation, the scaffolds (n=3 for each experimental condition) were washed with a 0.05% Tween solution, and stained with BCIP/NBT for 20 min at room temperature. Finally, they were washed with 0.05% Tween and stored in PBS (without Ca^2+^ and Mg^2+^) prior to imaging. Alizarin Red S (ARS) staining was used to assess calcium deposition and therefore mineralization of the scaffolds. After fixation, the scaffolds were washed with deionized water and exposed to 2% (w/v) ARS for 1h at room temperature. They were then washed with deionized water to remove the excess ARS staining solution and stored in PBS (without Ca^2+^ and Mg^2+^) prior to imaging. Finally, all scaffolds were imaged using a 12-megapixel digital camera.

### 2.5. Mineralization analysis using scanning electron microscopy and energy-dispersive spectroscopy

Scaffolds (n=3 for each experimental condition) were fixed in 4% para-formaldehyde for 48h and dehydrated in increasing concentrations of ethanol (from 50% to 100%), as previously described (21). They were then dried using a critical point dryer and gold-coated to a final coating thickness of 5 nm. Scanning electron microscopy (SEM) images were acquired with a JEOL JSM-7500F FESEM scanning electron microscope (JEOL, Peabody, MA) at 2 kV. Energy-dispersive spectroscopy (EDS) was performed on three different areas of each scaffold surface for mineral aggregate composition analysis.

### 2.6. Young’s modulus measurements

The Young’s modulus of the scaffolds after culture in non-differentiation or differentiation medium (n=3 for each experimental condition) was calculated following a compression test using a custom-built uniaxial compression apparatus and compared to that of decellularized apple-derived cellulose scaffolds without MC3T3-E1 seeded cells. The force and position were recorded with a 1.5 N load cell (Honeywell, Charlotte, NC) and an optical ruler, respectively. The force-displacement curves were obtained by compressing the samples (after removing them from the culture medium) at a constant rate of 3 mm min^-1^ and a maximum compressive strain of 10% of the construct height. The Young’s modulus was obtained by fitting the linear portion of the stress-strain curve between 9% and 10% strain.

### 2.7. Rat calvarial defect model

Experimental protocols were reviewed and approved by the Animal Care and Use Committee of the University of Ottawa. Bilateral craniotomy was performed following an established protocol (22). Male Sprague-Dawley rats (n=6) were anaesthetized with isoflurane, first at 3% until they were unconscious, and then at 2-3% throughout the procedure. After removing the periosteum, defects (5-mm diameter) were created in both parietal bones on each side of the sagittal suture using a dental drill equipped with a 5-mm diameter trephine, under constant irrigation of 0.9% NaCl. The surrounding bone was gently cleaned with 0.9% NaCl to remove any bone fragments. Decellularized scaffolds were prepared as described above, were cut into circular disks (5-mm diameter) with a biopsy punch and were placed in the 5-mm defects. The overlying skin was closed with sutures. Rats were given unlimited access to food and water and were monitored daily by certified animal technicians at the Animal Care & Veterinary Service of the University of Ottawa. Rats were euthanized by CO_2_ inhalation and thoracic perforation as secondary euthanasia measure, after 8 weeks post-implantation. The skin covering the skull was removed using a scalpel blade, exposing the cranium. The skull was cut at the frontal and occipital bones and on the side of both parietal bones using a dental drill, thereby completely removing the top section of the skull that was then processed for either mechanical assessment or histological analysis.

### 2.8. Push-out test

To assess the force required to push out the scaffolds from the surrounding bone, push-out tests were carried out after the 8 weeks of implantation using a uniaxial compression device (MTI Instruments, Albany, NY) and a 445N load cell (Omega Engineering, Norwalk, CT). The samples (n=7 from 4 animals) were placed on the sample holder of the instrument so that the dorsal side of the bone was facing up (Figure 4C). A plunger was slowly lowered at 0.5 mm/min until slightly touching the sample. The force vs. distance were recorded until complete push-out of the scaffolds, and the maximum force was recorded at the break point on the force vs. distance curve (Figure 4D).

### 2.9. Cell infiltration and mineralization analysis by histology

*In vitro* scaffolds (n=1 in non-differentiation medium and n=2 in differentiation medium) were fixed in 4% para-formaldehyde for 48h and stored in 70% ethanol before paraffin embedding, sectioning, and staining by the PALM Histology Core Facility of the University of Ottawa. Briefly, 5 μm-thick serial sections were stained with hematoxylin and eosin (H&E; ThermoFisher) or Von Kossa (VK; ThermoFisher), starting 1 mm inside the scaffolds. Slides were imaged using a Zeiss AXIOVERT 40 CFL microscope (Zeiss, Toronto, ON) to evaluate cell infiltration (H&E) and mineralization (VK). Image analysis was performed using ImageJ software.

*In vivo* scaffolds (n=4 from 2 animals) were fixed in 10% formalin (Sigma-Aldrich) for 72h and stored in 70% ethanol (Sigma-Aldrich), and all subsequent embedding, sectioning and staining was performed by AccelLAB Inc. (Boisbriand, QC). The scaffolds were embedded in methyl methacrylate and serially cut in 6-μm thick sections, at three different levels (top, bottom, and towards the center). Sections were stained with either H&E or Goldner’s Trichrome (GTC). Histological slides were imaged using a Zeiss AXIOVERT 40 CFL microscope to evaluate cell infiltration (H&E) and collagen deposition (GTC). Images were analyzed using ImageJ software.

### 2.10. Statistical analysis

All data are reported as mean ± standard error of the mean (S.E.M.). The data were assumed to be normally distributed. The Levene’s test was used to confirm that the data conformed to the assumption of homogeneity of variance. Statistical analysis was then performed using a one-way ANOVA followed by Tukey post-hoc tests for Young’s moduli mean comparison. A value of *p* < 0.05 was considered to be statistically significant.

## 3. RESULTS

### 3.1. Pore size measurement, cell distribution, and in vitro mineralization

Complete removal of native cellular components of the apple tissue was achieved after SDS and CaCl_2_ treatments (Figure 1A). The highly porous nature of the scaffolds was observed using confocal microscopy. Image quantification revealed an average pore size of 154 ± 40 μm. The pore size distribution ranged from 73 μm to 288 μm, with the majority of the pores ranging between 100 and 200 μm (Figure 1B).

**Figure 1.**
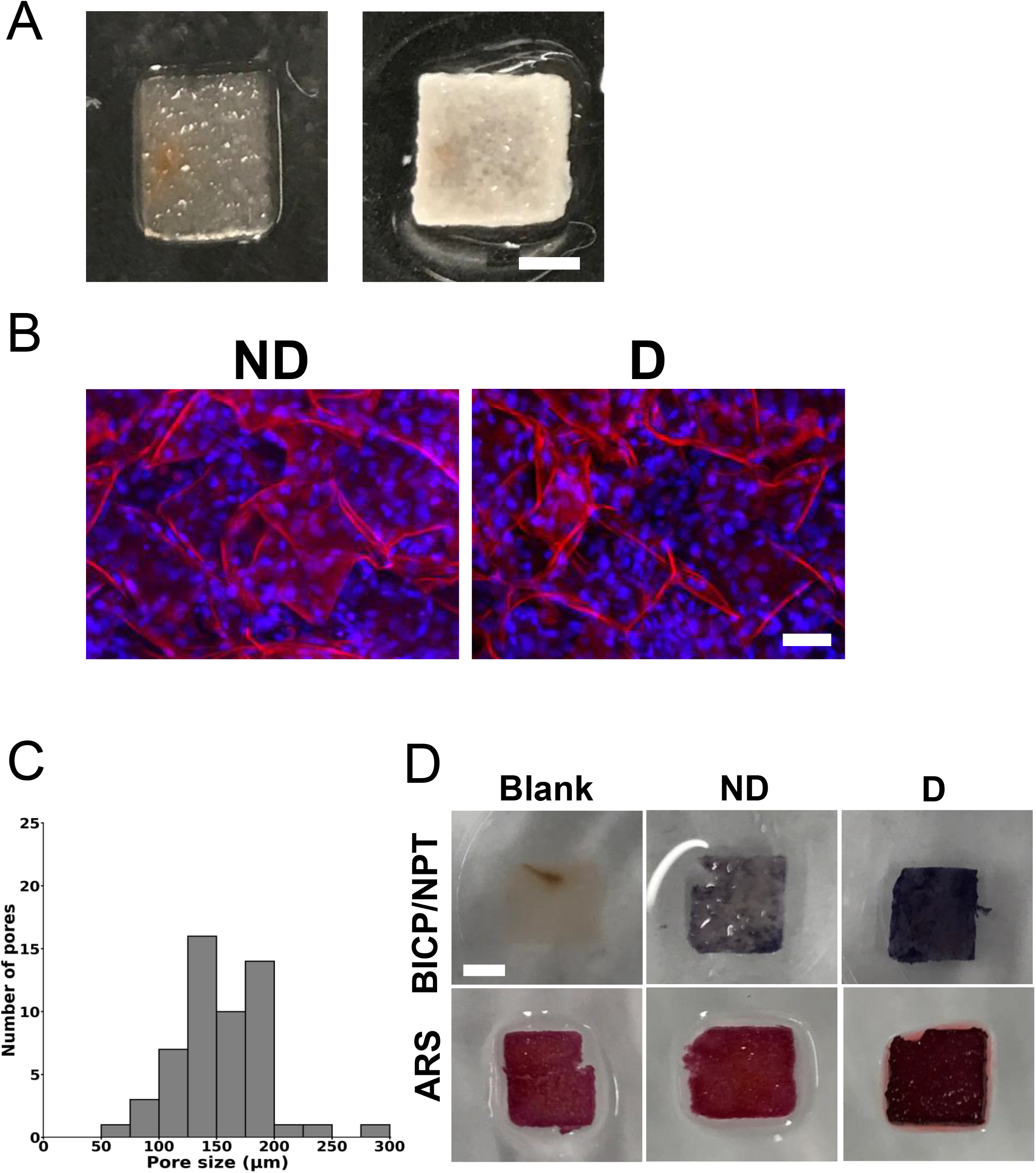
(A) Representative photographs of an apple-derived cellulose scaffold after removal of the plant cells and surfactant (left) and a MC3T3-E1-seeded scaffold after 4 weeks in osteogenic differentiation medium (right) (scale bar = 2 mm); (B) Representative confocal laser scanning microscope images showing seeded cells in a scaffold after 4 weeks of culture in non-differentiation medium (“ND”) or osteogenic differentiation medium (“D”) (scale bar = 50 μm). The scaffolds were stained for cellulose (red) and for cell nuclei (blue) using propidium iodide and DAPI staining, respectively; (C) Pore size distribution of decellularized apple-derived cellulose scaffolds before MC3T3-E1 cell seeding, from maximum projections in the Z axis of confocal images. A total of 54 pores were analyzed in 3 different scaffolds (6 pores in 3 randomly selected areas per scaffold); (D) Representative photographs of scaffolds stained with 5-bromo-4-chloro-3′-indolyphosphate and nitro-blue tetrazolium (BCIP/NBT) for alkaline phosphatase (ALP) activity or Alizarin Red S (ARS) for calcium deposition, to visualize mineralization (scale bar = 2 mm - applies to all). The scaffolds without cells (“Blank”) did not stain with BCIP/NBT. Stronger ALP activity was visualized by stronger blue contrast in the scaffolds cultured in differentiation medium (“D”), compared to their counterparts cultured in non-differentiation medium (“ND”). For the ARS staining, the blank scaffolds and the scaffolds cultured in non-differentiation medium (“ND”) displayed a lighter red color than the scaffolds cultured in differentiation medium (“D”). The calcium deposition was highlighted by a strong, dark red color in the scaffolds cultured in differentiation medium (“D”). Three different scaffolds (n=3) were analyzed for each experimental condition.

After 4 weeks of culture in differentiation medium, white mineral deposits were observed throughout the cell-seeded scaffolds (Figure 1A). Scaffolds with cells had a distinct opaque white colour, suggesting mineralization, that was not observed in the blank scaffolds (scaffolds without cells). Finally, confocal laser scanning microscopy analysis also showed that cells were homogenously distributed in the scaffolds (Figure 1A).

To analyze ALP activity and mineralization, the scaffolds were stained with BCIP/NBT and ARS, respectively (Figure 1C). The BCIP/NBT staining revealed that ALP activity increased significantly (as indicated by the strong purple colour) in cell-seeded scaffolds cultured in differentiation medium compared to blank scaffolds or cell-seeded scaffolds cultured in non-differentiation medium. Similarly, cell-seeded scaffolds cultured in differentiation medium displayed a stronger red color after ARS staining, indicating a higher degree of calcium mineralization, than the blank scaffolds or the cell-seeded scaffolds cultured in non-differentiation medium. Some background staining was clearly visible in the blank scaffolds, possibly due to the use of CaCl_2_ in the decellularization protocol.

Staining (H&E and VK) as well as SEM and EDS were used to analyze cell infiltration and further evaluate mineralization. H&E staining (Figure 2A) showed that the cell-seeded scaffolds cultured in either non-differentiation or differentiation medium displayed good cell infiltration, with multiple nuclei visible in the periphery and through the scaffolds. Collagen was also visible in pale pink. In addition, VK staining revealed that the pore walls of the scaffolds were stained after 4-weeks of culture in differentiation medium. The pore walls of the scaffolds cultured in non-differentiation medium only showed the presence of calcium deposition on the outside periphery of the constructs, likely because of the absorption of calcium from the decellularization treatment. SEM analysis of the cell-seeded scaffolds (Figure 2B) revealed localized mineralization on the cell-seeded scaffolds cultured for 4 weeks in differentiation medium. Mineral deposits appeared as globular aggregates on the edge of the pores. No mineral aggregates were visible on the cell-seeded scaffolds cultured for 4 weeks in non-differentiation medium and on blank scaffolds. EDS spectra acquired on selected regions of interest, namely on the mineral aggregates on the cell-seeded scaffolds and on pore walls of the blank scaffolds, clearly displayed distinct characteristic signals corresponding to the deposition of phosphorous (P) and calcium (Ca) on the cell-seeded scaffolds cultured for 4 weeks in differentiation medium (Figure 2B).

**Figure 2.**
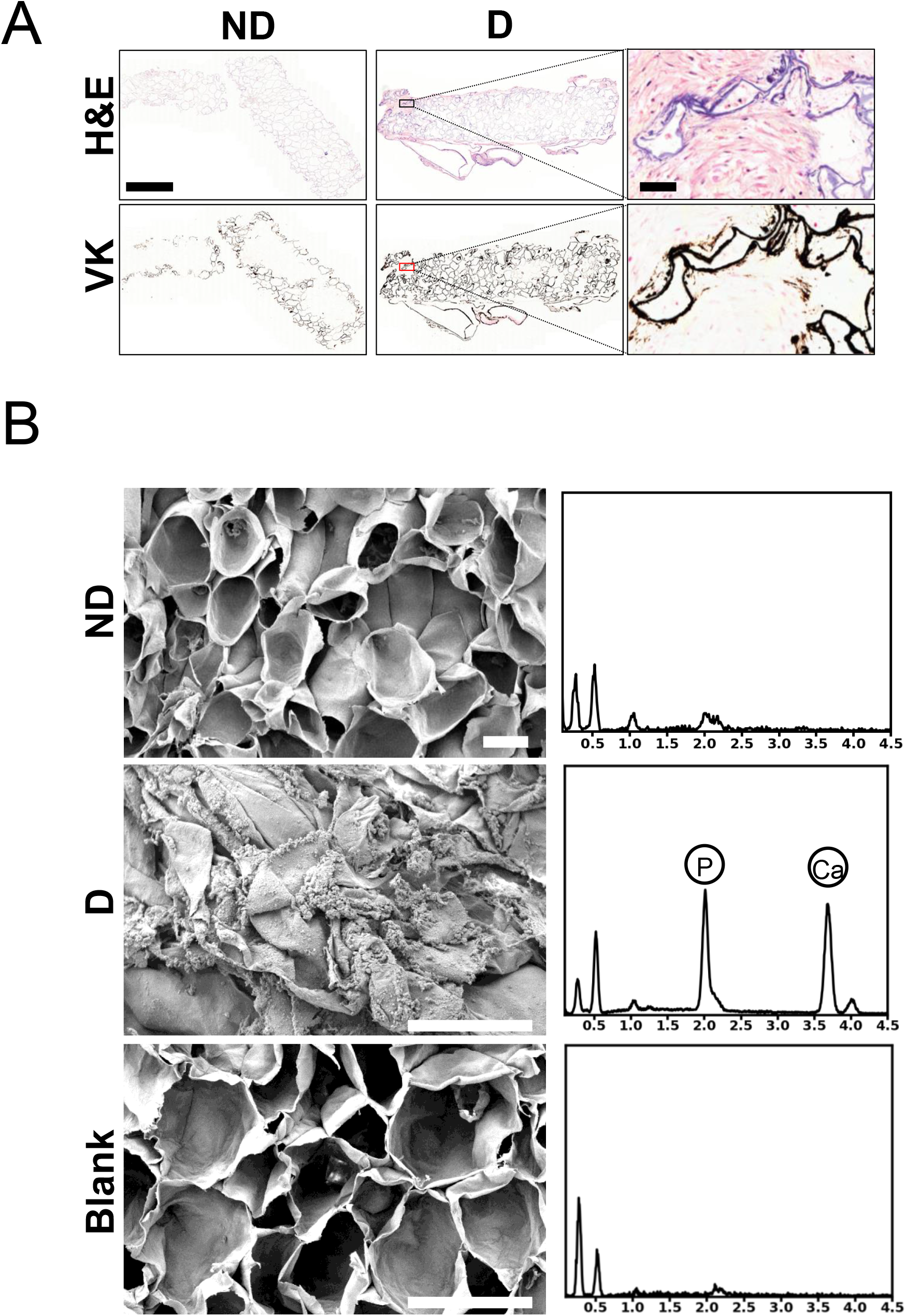
(A) Representative images of scaffold histological cross-sections. Paraffin-embedded scaffolds were cut into 5 μm-thick sections and stained with Hematoxylin and Eosin (H&E) to visualize cell infiltration or Von Kossa (VK) to visualize mineralization. Scaffolds were infiltrated with MC3T3-E1 cells with multiple nuclei and cytoplasm visible at the periphery and throughout the scaffolds (blue and pink, respectively). Collagen was also visible in pale pink. The pore walls in the scaffolds cultured in non-differentiation medium (“ND”) only showed the presence of mineralization at their periphery. The pore walls in the scaffolds cultured in differentiation medium (“D”) were entirely stained in black. The analysis was performed on one scaffold cultured in non-differentiation medium (“ND”) and on 2 scaffolds cultured in differentiation medium (“D”) (Scale bar = 1 mm for the lower magnification pictures and 50 μm for the higher magnification pictures); (B) Representative scanning electron microscopy micrographs and energy-dispersive spectra: Scaffolds were gold-coated and imaged using a JEOL JSM-7500F FESEM scanning electron microscope at 3.0 kV (scale bar= 100 μm - applies to all). Energy-dispersive spectroscopy spectra were acquired on each scaffold. Phosphorus (2.013 keV) and calcium (3.69 keV) peaks are indicated on each spectrum. Three different scaffolds were analyzed for each condition. Blank: scaffolds without seeded cells.

### 3.2. In vitro biomechanical analysis

The Young’s modulus of cell-seeded scaffolds was measured after 4 weeks of incubation in either non-differentiation or differentiation medium and compared to that of the blank scaffolds without cells) (Figure 3). Results showed no significant difference in the Young’s modulus between the blank scaffolds (31.6 ± 4.8 kPa) and the scaffolds cultured in non-differentiation medium (24.1 ± 8.8 kPa; *p*=0.88). On the other hand, a significant difference was observed between the blank scaffolds (31.6 ± 4.8 kPa) and the scaffolds cultured in differentiation medium (192.0 ± 16.6; *p*<0.001). Furthermore, the Young’s moduli of the cell-seeded scaffolds cultured in non-differentiation and differentiation medium were also significantly different (*p<*0.001).

**Figure 3.**
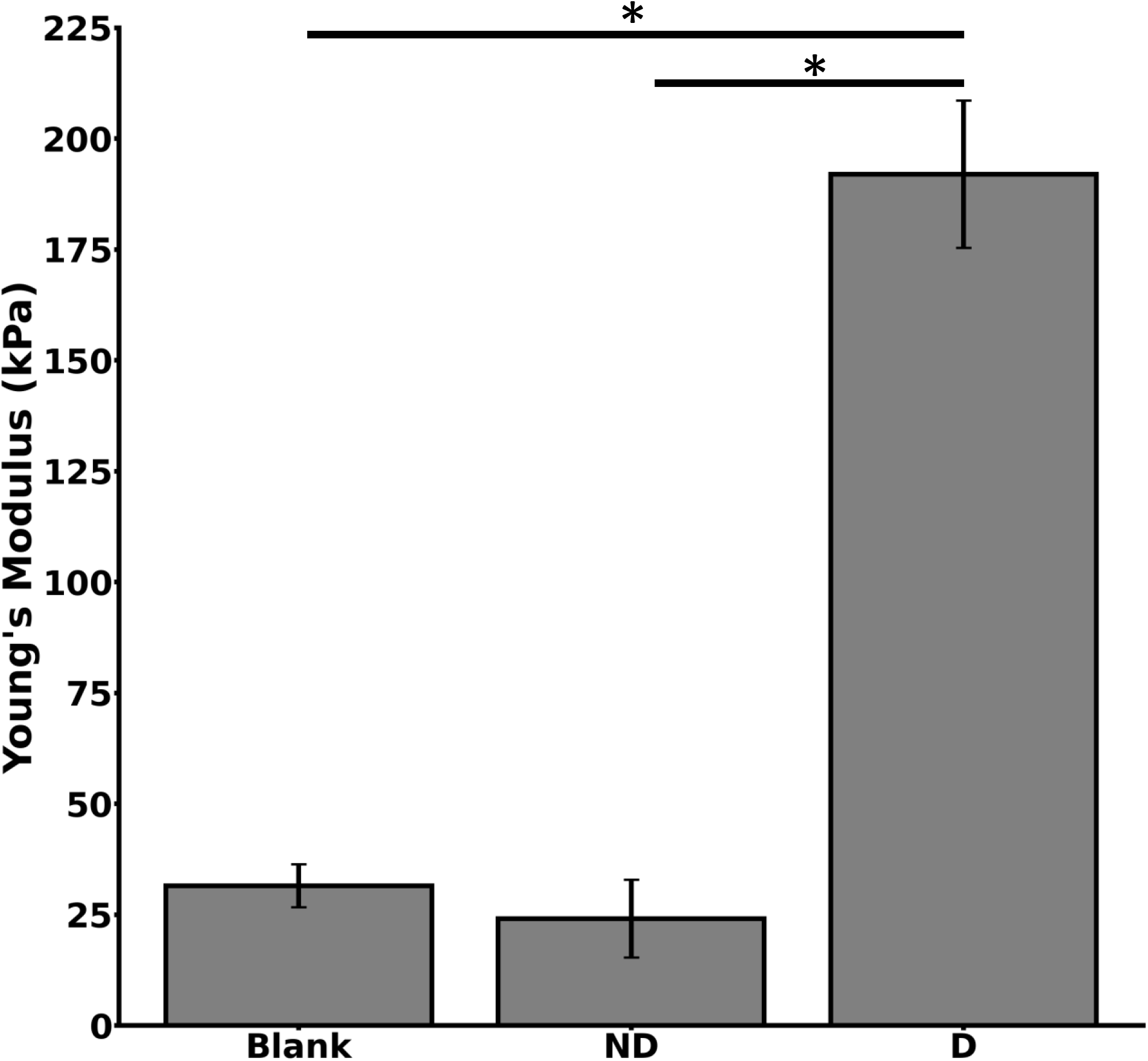
Young’s modulus of scaffolds without seeded cells (“Blank”) and of cell-seeded scaffolds after 4 weeks of culture in either non-differentiation (“ND”) or differentiation (“D”) medium. Statistical significance (* indicates *p*<0.05) was determined using a one-way ANOVA and Tukey post-hoc tests. Data are presented as mean ± S.E.M. of three replicate samples for each condition.

### 3.3. *In vivo* bone regeneration and biomechanical performance

Craniotomies were performed on Sprague-Dawley rats. Bilateral 5-mm diameter defects were created in both parietal bones, and apple-derived cellulose scaffolds (without seeded cells) were implanted in the defects (Figure 4A, B). The top section of the skull was retrieved and processed for either mechanical assessment or histology after 8 weeks.

**Figure 4.**
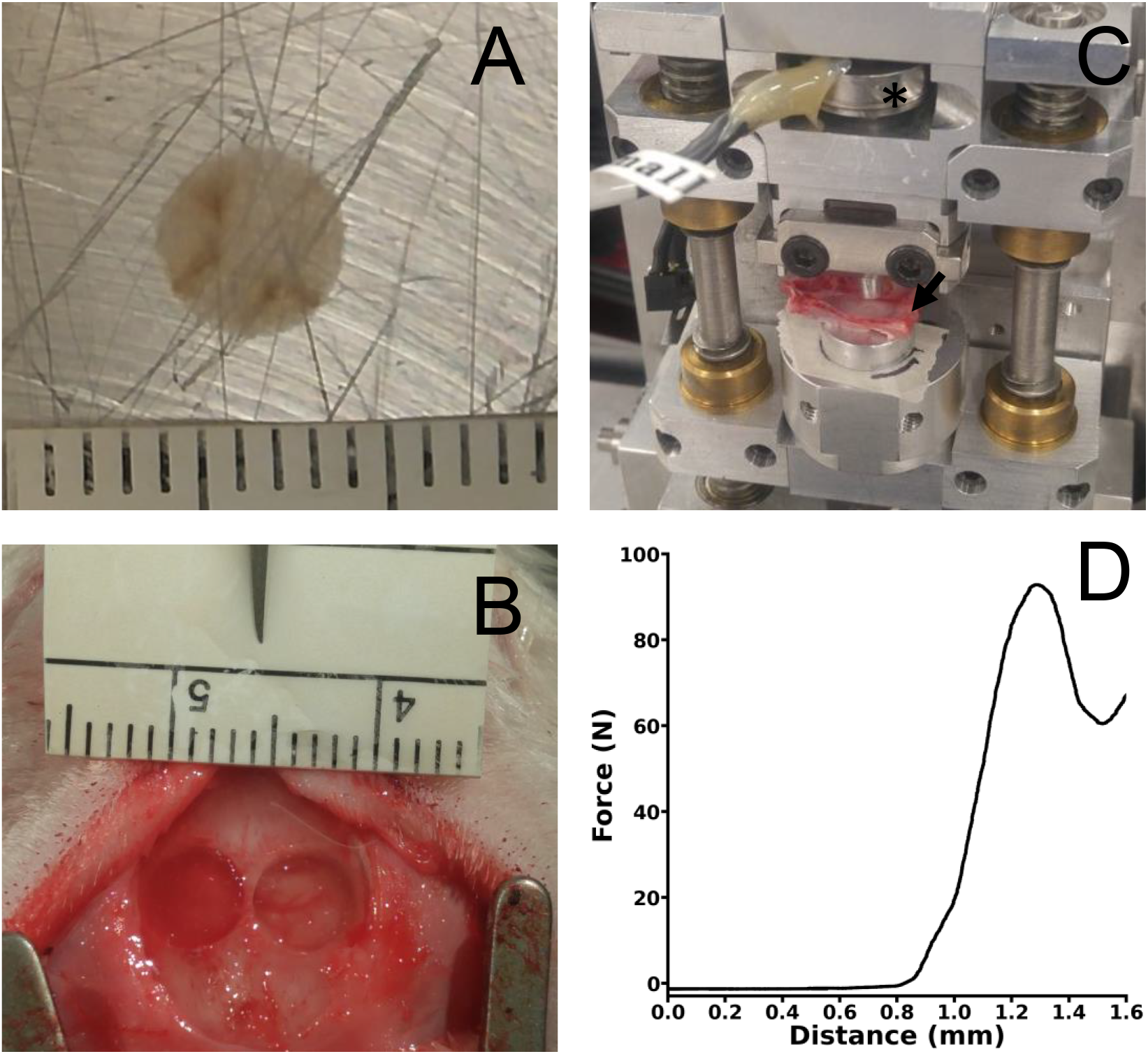
(A) Scaffolds; (B) Exposed skull with bilateral defects; (C) Photograph of uniaxial compression device for the push-out tests (the asterix (*) indicates the load cell; the arrow indicates the sample); (D) Typical force-displacement curve obtained during a push-out test.

Upon visual inspection, the scaffolds appeared to have been well integrated with the surrounding tissues of the skull. Mechanical push-out tests were performed to quantitatively assess the integration. Measurements were performed using a uniaxial compression device (Figure 4C) immediately after euthanasia of the animals. Results revealed that the average force required to dislodge the scaffolds from the surrounding bone was 113.6 ± 18.2 N.

Finally, histological analysis was performed to evaluate cell infiltration and extracellular matrix deposition within the grafted scaffolds after 8 weeks of implantation (Figure 5). H&E staining showed infiltration of cells within the pores of the scaffolds. Evidence of vascularization (as depicted by blood vessels) was also observed within the scaffolds and GTC staining showed the presence of type 1 collagen within the scaffolds.

**Figure 5.**
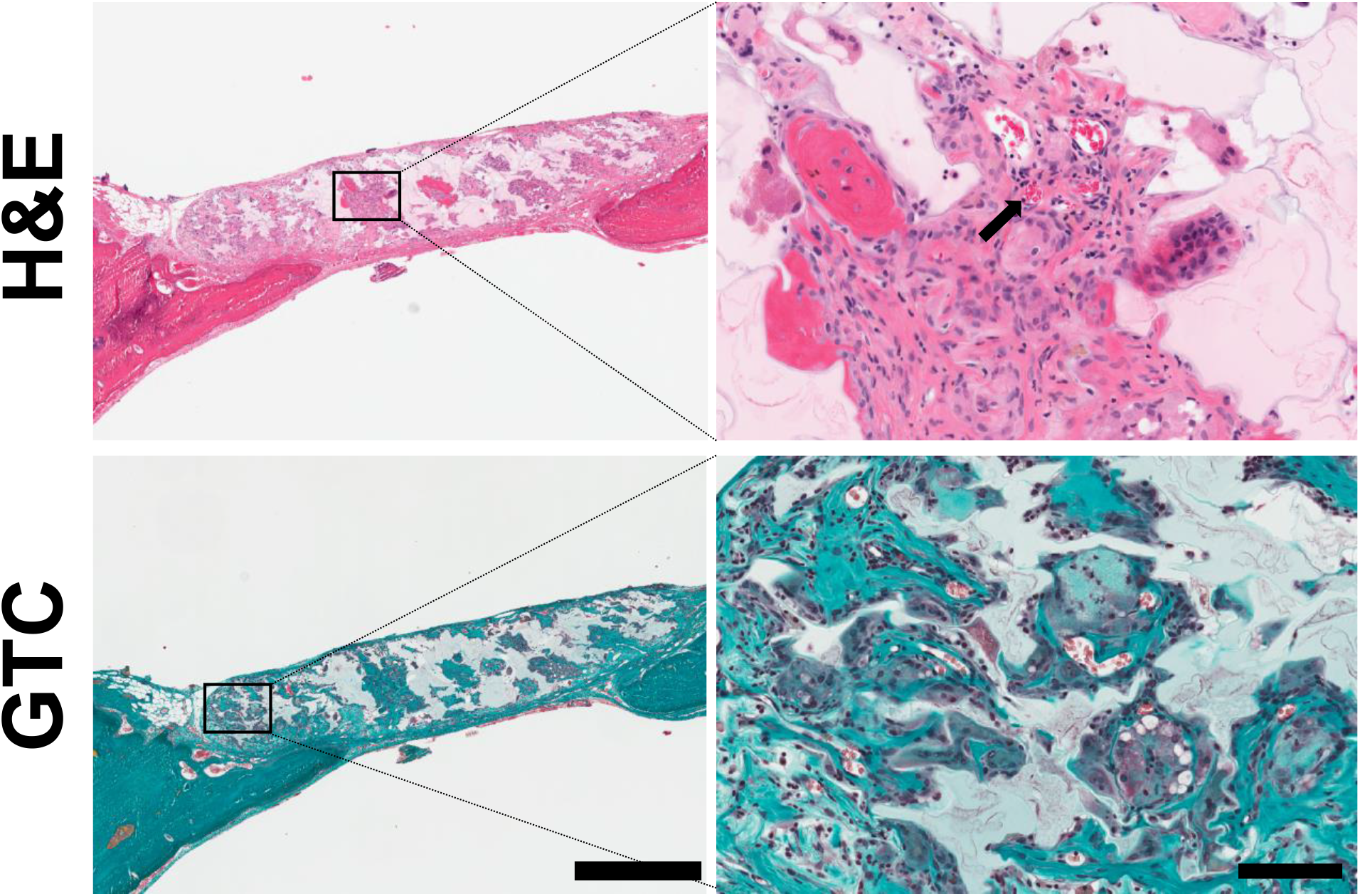
Representative images of implanted non-seeded scaffolds histological cross-sections after 8 weeks. Sections were stained with either hematoxylin and eosin (H&E) to visualize cells or Goldner’s Trichrome (GTC) to visualize type-1 collagen. The arrow indicates red blood cells. Presence of collagen is visible (scale bar = 1 mm and 200 μm for the left and right insets, respectively).

## 4. DISCUSSION AND CONCLUSIONS

*In vitro* and *in vivo* studies have shown the biocompatibility of plant-derived cellulose and their potential use for tissue engineering (14–18). Moreover, their uses for hosting osteogenic differentiation has been demonstrated (19). The aim of the present study was to further examine the potential of apple-derived cellulose scaffolds for BTE applications and analyze their mechanical properties *in vitro* and *in vivo*.

For *in vitro* studies, pre-osteoblast cells (MC3T3-E1) were seeded in the scaffolds after removing the native cells from the apple tissue, and were induced to undergo osteogenic differentiation. Results showed that the cells were able to proliferate and differentiate within the scaffolds, thereby demonstrating the potential for plant-derived cellulose scaffolds to support bone formation. A large number of cell nuclei were observed throughout the scaffold pores, similarly to the observations reported in previous studies (14–16,19). Moreover, similarly to our previous findings (14) and those of another group (19), we observed that the average diameter of the scaffold pores was ~154 μm, with the majority of the pores between 100 and 200 μm (Figure 1B). These sizes are consistent with the optimum pore size for bone growth, which has been shown to be in the range of 100–200 μm (7).

Staining analysis revealed a higher expression of ALP and the presence of more calcium deposits on the surface of the cell-seeded scaffolds after 4 weeks of culture in differentiation medium than in the blank scaffolds (without seeded cells) and in the cell-seeded scaffolds cultured in non-differentiation medium. Importantly, similar results with differentiated hiPSCs in apple-derived scaffolds have been observed (19). Histological analysis further confirmed that the constructs were mineralized by the infiltrated osteoblasts after differentiation. Of note is that the periphery of the constructs cultured in non-differentiation medium was also stained with VK. This non-specific staining may have been due to residual CaCl_2_ in the scaffolds after the decellularization process. Mineralization was further assessed by qualitative analysis of SEM pictures. After culture in differentiation medium, cell-seeded scaffolds displayed signs of ECM mineralization, with aggregates of minerals visible on the scaffold surface, specifically at the edges of the pores, which is consistent with previous studies using ECM scaffolds (23) and plant scaffolds (19). These aggregates were not visible on the surface of the scaffolds without cells. EDS analysis of the aggregates revealed high level of P and Ca, thereby suggesting the presence of apatite.

A significant change (approximately 8 fold increase) in the Young’s modulus of the scaffolds was demonstrated after culture in differentiation medium. On the other hand, the modulus of the scaffolds cultured in non-differentiation medium was similar to that of the blank scaffolds (without seeded cells). However, It should be noted that despite the increase in the Young’s modulus of the scaffolds cultured in differentiation medium, the moduli remained much lower than that of bone (0.1 to 2 GPa for trabecular bone and 15 to 20 GPa for cortical bone) (8), cancellous allograft (3.78 GPa) (24) alloplastic grafts made of poly-ether-ether-ketone (3.84 GPa) (24), titanium (50.20 GPa) (24), and cobalt-chromium alloy (53.15 GPa) (24) implants. Disparity between the Young’s modulus of the scaffolds and the surrounding bone can cause stress shielding (25). The imbalance of stress distribution at the bone-implant interface can thus lead to bone augmentation or bone resorption around the implant and ultimately implant failure (25). Therefore, in the current formulation, these scaffolds may not be appropriate for load-bearing applications.

Decellularized apple scaffolds were then implanted in 5 mm critical-sized cranial defects in rats. Mechanical assessment indicated an average force of 113.6 ± 18.2 N to dislodge the scaffolds from the surrounding bone. This force is similar to the force required to displace intact calvarial bone (127.1 ± 9.6◻N) (20). This indicates that the scaffolds integrated well to the surrounding bone and connective tissues. Moreover, the force is similar to the one reported after 8 weeks implantation of calcium-deficient hydroxyapatite scaffolds loaded with bone morphogenic protein-2 (119.1 ± 17.8◻N) (20).

In summary, this study confirmed that pre-osteoblasts can adhere and proliferate within apple-derived cellulose scaffold constructs. Mineralization occurred within cell-seeded scaffolds after chemically inducing osteogenic differentiation of pre-seeded pre-osteoblasts, which resulted in a significant increase in the Young’s modulus of the constructs, although it remained much lower than that of natural bone. Moreover, the force required to dislodge the implanted plant-derived cellulose scaffolds was similar to the one observed with calvarial bone and other type of scaffolds used for BTE. Similarly to previous reports, cells infiltrated the scaffolds and deposited type-1 collagen. Overall, these results show that plant-derived cellulose scaffolds have potential for BTE applications, which is consistent with the findings of a previous study from a different group (19). However, the difference in stiffness compared to trabecular or cortical bone will likely require the development of composite biomaterials to better match the mechanical properties of living bone. While interest in the use of plant-derived scaffolds for BTE has grown in recent years, this biomechanical mismatch limits their applications, especially for load-bearing conditions. Re-engineering plant-derived cellulose scaffolds through chemical modification or creating composites with other biological/synthetic polymers, as we have previously shown (16), may be required for load bearing applications.

## ACKNOWLEDGMENTS

This work was supported by a Discovery Grant from the Natural Sciences and Engineering Research Council of Canada (NSERC) and a grant from the Li Ka Shing Foundation. M.L.L. was supported by the Ontario Centers of Excellence TalentEdge program. R.J.H. was supported by an NSERC postgraduate scholarship and an Ontario Graduate Scholarship (OGS).

